# Cryo-EM structure of the NLRP3 decamer bound to the cytokine release inhibitory drug CRID3

**DOI:** 10.1101/2021.07.22.453353

**Authors:** Inga V. Hochheiser, Michael Pilsl, Gregor Hagelueken, Jonas Moecking, Michael Marleaux, Rebecca Brinkschulte, Eicke Latz, Christoph Engel, Matthias Geyer

**Affiliations:** Institute of Structural Biology, University of Bonn, Venusberg-Campus 1, 53127 Bonn, Germany; Structural Biochemistry Group, Regensburg Center for Biochemistry, University of Regensburg, Regensburg, Germany; Institute of Innate Immunity, University of Bonn, Venusberg-Campus 1, 53127 Bonn, Germany

**Keywords:** NLRP3, CP-456.773, CRID3, MCC950, cryo-EM, AAA+ ATPase, NOD-like receptors, inflammasome

## Abstract

NLRP3 is an intracellular sensor protein whose activation by a broad spectrum of exogenous and endogenous stimuli leads to inflammasome formation and pyroptosis. The mechanisms leading to NLRP3 activation and the way how antagonistic small molecules function remain poorly understood. Here we report the cryo-electron microscopy structures of full-length NLRP3 in its native form and complexed with the inhibitor CRID3 (also named MCC950). Inactive, ADP-bound NLRP3 is a decamer composed of homodimers of intertwined LRR domains that assemble back-to-back as pentamers with the NACHT domain located at the apical axis of this spherical structure. Molecular contacts between the concave sites of two opposing LRRs are mediated by an acidic loop extending from an LRR transition segment. Binding of CRID3 significantly stabilizes the NACHT and LRR domains relative to each other, allowing structural resolution of 3.9-4.2 Å. CRID3 binds into a cleft, connecting four subdomains of the NACHT with the transition LRR. Its central sulfonylurea group interacts with the Walker A motif of the NLRP3 nucleotide-binding domain and is sandwiched between two arginines from opposing sites, explaining the specificity of NLRP3 for this chemical entity. With the determination of the binding site of this lead therapeutic, specific targeting of NLRP3 for the treatment of autoinflammatory and autoimmune diseases and rational drug optimization are within reach.

---

Inflammasomes are cytosolic sensor proteins that recognize endogenous damage signals and exogenous microbial invasions^1^. Upon activation, the proteins of the family of nucleotide-binding oligomerization domain (NOD)-like receptors (NLRs) form supramolecular complexes that lead to caspase maturation, gasdermin D cleavage, the release of cytokines, and ultimately pyroptotic cell death^2^. NLRP3 is a well-studied member of the NLRs that senses cellular stress caused by bacterial, viral and fungal infections, as well as sterile inflammation^3^. While much has been learned about the triggers of NLRP3, the activation mechanism remains obscure.

NLRP3 is composed of an N-terminal Pyrin domain (PYD) that is the effector domain for supramolecular complex formation; a central triple-ATPase domain called NACHT that mediates protein oligomerization upon activation; and a C-terminal leucine-rich repeat (LRR) domain that may act as the signal sensor^2^. Basal, ADP-bound NLRP3 is thought to be in an autoinhibited state that is relieved by the action of various kinases. This includes phosphorylation of the activation loop upstream of the nucleotide-binding domain (NBD) and interaction with the mitotic Ser/Thr kinase NEK7^4-7^. Activation, concomitant with ATP-binding in the NBD, is leading to a conformational rearrangement of the NACHT subdomains helical domain 1 (HD1), winged helix domain (WHD), and helical domain 2 (HD2). Multiple autosomal mutations of the *NLRP3* gene have been identified that lead to its aberrant activation in systemic and joint inflammation described as cryopyrin-associated periodic syndrome (CAPS) and complex diseases such as multiple sclerosis, type 2 diabetes, Alzheimer’s disease and atherosclerosis^8^. However, how endogenous NLRP3 resides in the inactive state is currently unknown.

The cytokine release inhibitory drug 3 (CRID3) belongs to the sulfonylureas and has been first described in 2003 (named CP-456.773) resulting from an optimization screen of the anti-diabetic drug glibenclamide^9-11^. While inflammasomes were not known at this time, CRID3 has been shown to have strong anti-inflammatory efficacy inhibiting IL-1β processing. Later, its target protein was identified as NLRP3, showing that CRID3 potently inhibits ATP-triggered, NLRP3-mediated IL-1β release with an IC_50_ value of 8.1 nM in human monocytes^12-14^. In this study, we determine the cryo-EM structures of full-length human NLRP3 in an inactive state and bound to the antagonist CRID3.

## Full-length NLRP3 forms a defined multimer

Recombinant full-length, wild type NLRP3 (residues 3-1036) was expressed in baculovirus infected insect cells as MBP fusion protein and purified to homogeneity. The protein elutes in two peaks in size exclusion chromatography experiments, one close to the void volume indicative of a high molecular weight assembly, and one at a defined size larger than 1 MDa (Extended Data Figs. 1a,b). Both peaks could be separated by fractionation and remained stable, indicating that these two conformers are not in a concentration-dependent dynamic equilibrium. SEC-MALS experiments revealed an average molecular mass of 1.58 MDa corresponding to 10-times the mass of the MBP-NLRP3 protein and cleavage of the MBP-tag retained the oligomeric assembly of Peak 2 (Extended Data Figs. 1c-e).

Negative stain electron microscopy images of untagged, full-length NLRP3 showed homogenous particles of circular shape with a diameter of 20 nm in accordance with a 1.2 MDa protein complex. A cryo-EM dataset of BS3-crosslinked NLRP3 containing 9,186 movies was recorded using a Titan Krios microscope equipped with a K3 detector. Image analysis showed average class sums with a two-fold and five-fold rotational symmetry. 3D reconstruction at a resolution of 8-11 Å revealed an overall spherical structure with a pentameric assembly at the polar sites and a meander ring along the equator (Extended Data Fig. 2). Using the NACHT and LRR domains from the NLRP3–NEK7 complex^15^ for model building, it became apparent that the LRRs are interlocked with their concave sites along the horizontal plane. The NACHT domains instead assemble to a pentamer at the vertical axis.

To increase the particle stability, we added the molecular inhibitor CRID3 to the expression and purification procedure. Already from SEC runs and negative stain imaging, it was evident that CRID3 significantly stabilizes the conformation of NLRP3 (Extended Data Fig. 1e,f). A large cryo-EM dataset of 14,336 movies was collected on a Titan Krios equipped with K3 direct electron detector initially yielding 1,588,061 particles. Processing resulted in a 3.9-4.2 Å cryo-EM map reconstructed from 379,240 particles selected following multiple rounds of 2D and 3D classifications (Methods). The density for NLRP3 was improved by focused classifications and masked refinements, including NLRP3 dimer or trimer assemblies and either the LRR or NACHT domains alone (Extended Data Figs. 3, 4). This led to a reconstruction of the NLRP3 decamer at an overall resolution of 4.1 Å, with a local resolution of around 3.9 Å for the LRR and the NACHT module (Fig. 1). Initial model building was performed with separated subdomains NBD, HD1 and WHD of the NACHT based on the NLRP3–NEK7 complex structure (6NPY)^15^, a model of a canonical LRR (based on 2BNH)^16^, whereas FISNA, HD2 and the transition LRR were built from models derived from the NOD2 structure (5IRN)^17^. Resulting models were validated by comparison to model-independent secondary structure predictions. No density was visible for the N-terminal PYD suggesting that this domain is flexibly linked to the NACHT domain (Extended Data Figs. 5, 6).

**Fig. 1.**
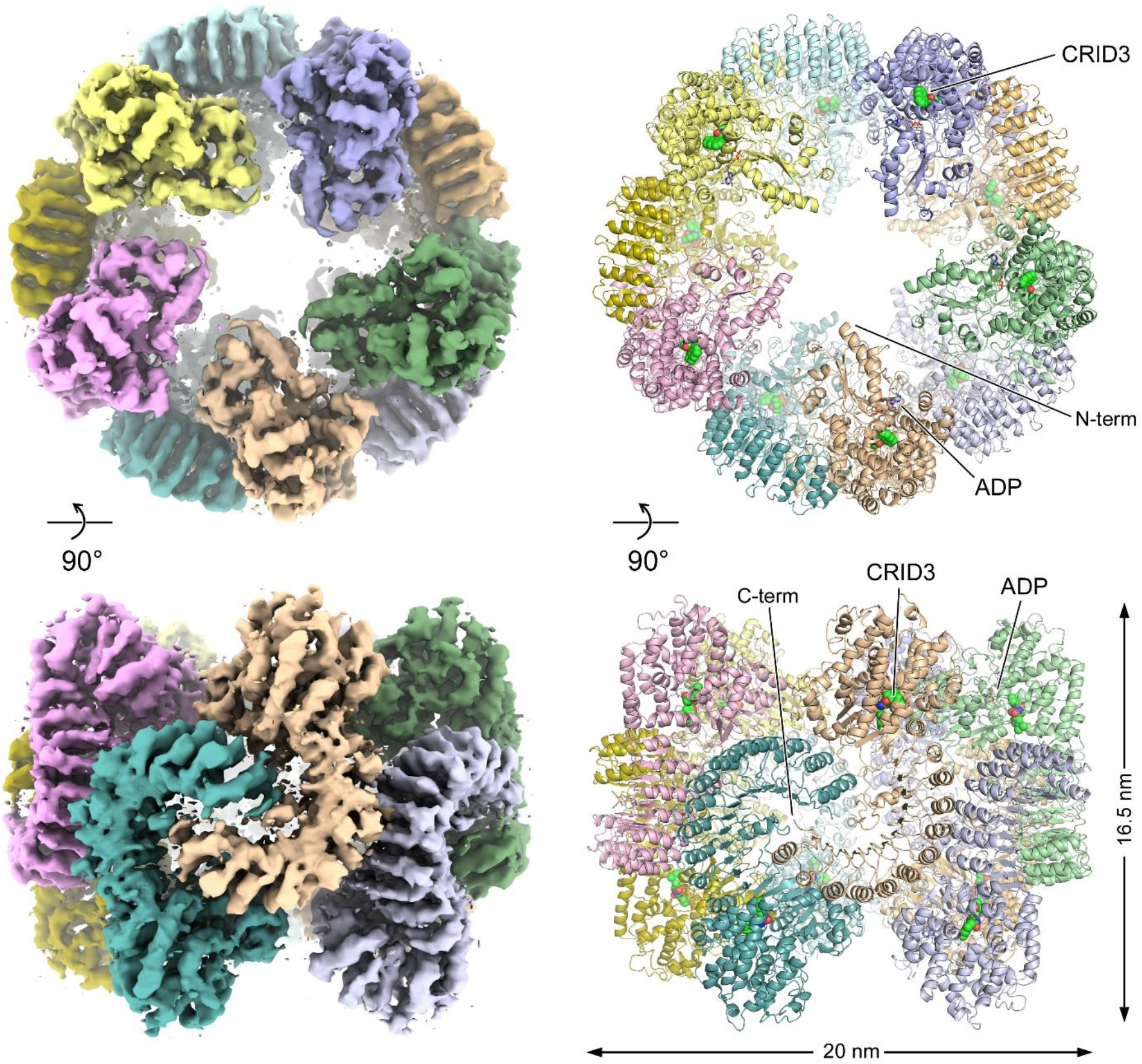
Structure of the NLRP3 decamer. Apical (top) and equatorial (bottom) view on the NLRP3 decamer. The cryo-EM density of a composite map from the decamer structure is shown at a threshold level of 0.0257 (left). Two views on the decamer structure with NLRP3 shown as ribbon (right). The NACHT domain is at the top with the intertwined LRRs forming a ring-like structure in the centre. ADP (grey) is shown in stick representation and CRID3 (green) as spheres.

## Structure of the NLRP3 decamer

NLRP3 consists of seven subdomains that we identify in the atomic model as FISNA (131-218), NBD (219-372), HD1 (373-434), WHD (435-541) and HD2 (542-651) for the NACHT domain and transition LRR (trLRR; 652-742) and canonical LRR (cnLRR; 743-1036) for the LRR domain (Fig. 2a). The atomic structure was built as a continuous chain from residues 133-1036, including ADP and CRID3, but with gaps in the density map for residues 153-159 and 181-197 (Fig. 2b). As seen before for NLRC4, NOD2, NAIP5 and NLRP3–NEK7 structures^15,17-19^, the four subdomains of the NACHT are tightly interconnected. The nucleotide-binding site of the NBD α/β roll is covered by the preceding FISNA through formation of an anti-parallel β-strand with the terminal fifth strand of the β-sheet characteristic for AAA+ ATPases^20^, precluding a free exchange of ATP/ADP. The LRR region consists of twelve helices on the outer side and 13 β-strands on the concave side. From position 743 on, the LRR follows a rigorous 28/29 amino acid alternation as seen, e.g., for the RNase inhibitor protein^16^. The sequence preceding this canonical LRR is highly variable in NLRs and includes an acidic loop 689-702 unique for NLRP3. Overall, the LRR in the NLRP3 decamer is in a partly open conformation, similarly to the NLRP3–NEK7 complex but different from the NLRC4 or NOD2 structures (Extended Data Fig. 7).

**Fig. 2.**
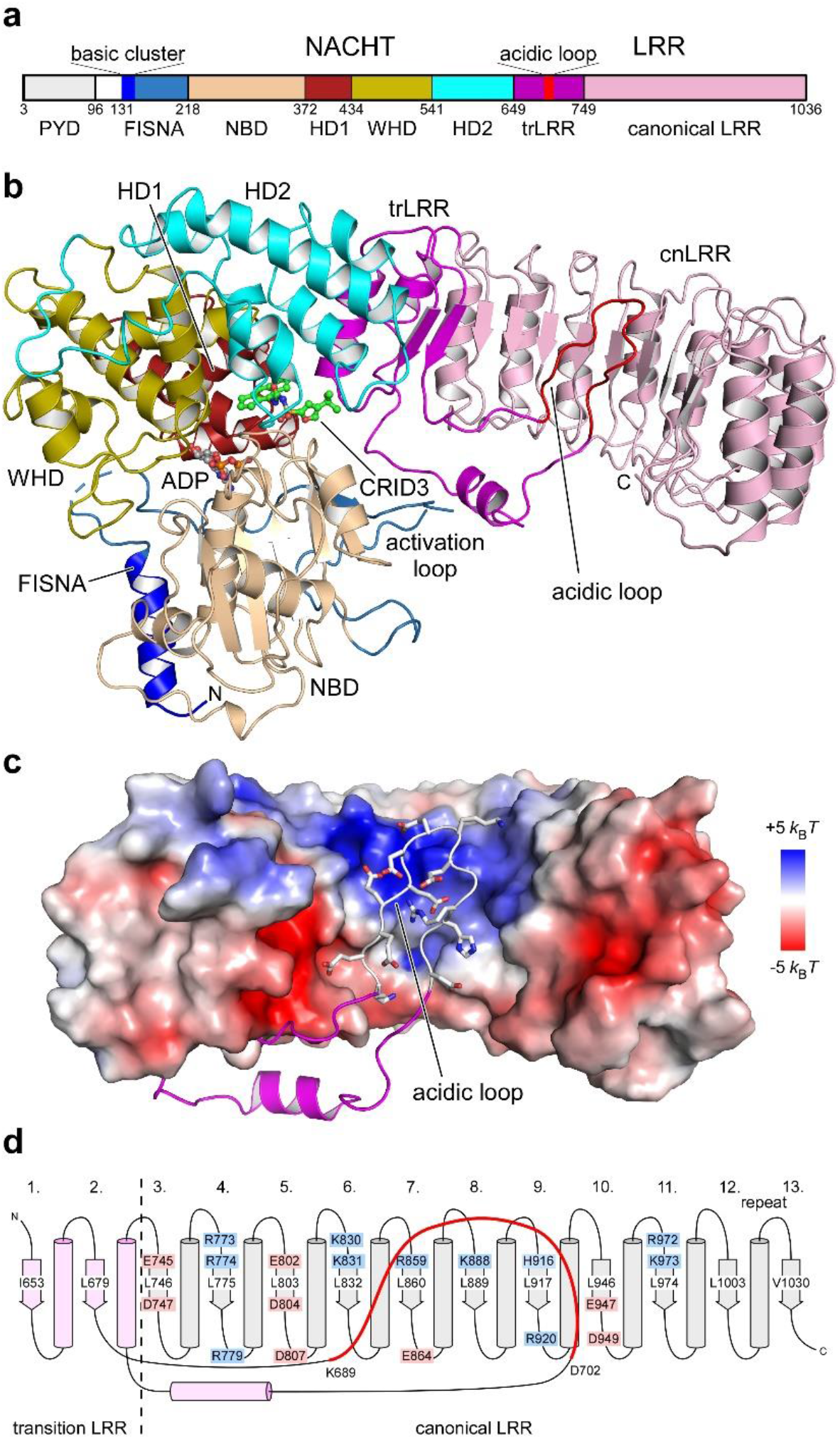
The acidic loop interacts with the concave site of the LRR. **a**, Domain composition of human NLRP3 with domain boundaries indicated. The basic cluster (131-147) and the acidic loop (689-702) are drawn as bars. **b**, Cartoon diagram of NLRP3 as determined from the decamer structure. Subdomains are colour-coded as in a, and ADP and CRID3 ligands labelled. **c**, Electrostatic surface potential of the LRR domain (650-1036) with the acidic loop drawn as chain. A positively charged rim is formed in the funnel of the concave β-strand structure that is covered by the acidic loop. **d**, Schematic of the LRR region with charged amino acids in the concave surface displayed. The acidic loop conformation (689-702) interacting with this surface is coloured red. The central leucine of the LxxLxL consensus motif in each repeat is indicated.

While the secondary structure elements and overall conformation of full-length NLRP3 are similar to the structure of NLRP3ΔPYD in the NLRP3–NEK7 complex^15^, there is a shift in the amino acid register ranging from -15 residues in the first helix of HD2 to +44 residues in the first β-strand of trLRR (Extended Data Fig. 8). This shift could be explained by a different arrangement of flexible loops in the decamer structure starting at position E538 in the linker region between WHD and HD2 and continuing along the entire length of the trLRR, including the acidic loop. It harmonizes in the α-helix of the second trLRR repeat at position L737, which marks the beginning of the canonical LRR domain. How NEK7 binding could induce this distinct shift in the most versatile and characteristic HD2 and trLRR region of NLRP3 will be analysed in future studies.

## An acidic loop binds to the concave LRR

The transition LRR starts in the decamer structure with the sequence F_650_xxIxI, forming the first β-strand of a 26-residue repeat. The following repeat loops after the β-strand into the concave side of the succeeding LRR. The electrostatic surface potential of the inner torus ring is highly charged with a basic rim at its upper site (Figs. 2c,d). The acidic loop sequence KEEEEEEKEGRHLD (689-702) reaches out to this basic surface in an antiparallel lobe, forming multiple electrostatic interactions with the LRR’s convex surface. The EEKE sequence is at the tip of this loop; it drops back to the bottom of the LRR and forms another helical twist before proceeding into an outer helix and completing the trLRR subdomain. This unexpected conformation is facilitated by several residue ‘mismatches’ that preclude a proper LRR fold and may induce the flexibility of the transition LRR. These mismatched residues are particularly phenylalanines 729 and 734 that reside at positions, otherwise occupied by glycine/serine or cysteine/serine residues, respectively (Extended Data Fig. 9). Likewise, the small A715 and polar H718 residues should instead be leucines for a bona fide LRR β-strand. The canonical LRR following the trLRR starts at position L743 and continues to the end of the protein with alternating 28/29 residue tandem repeats, each encoded by an individual exon^21^. The outer helix following the acidic loop is close to the FISNA activation loop with the two phosphorylation sites S_198_P and S_201_P.

## Intermolecular interfaces in the decamer

There are three types of unique interfaces between NLRP3 subunits in the decamer assembly, designated as *A, B* and *C* (Fig. 3a). In interface *A*, the LRRs of two molecules face each other with their concave sites, with the C-terminal end of one molecule engaging the groove of the opposing molecule (Fig. 3b). The interaction is mediated by residues of the last helix of the canonical LRR domain and the very C-terminus, contacting the acidic loop of the transition LRR. This interaction of interlaced LRRs is stabilized by polar contacts in interface *B*, where residues on the surface of the second last helix of the LRR interact with the loop sequence K_617_AKKLQ. This basic loop is located between the third and fourth helix of HD2 and extends with the two consecutive lysines at its tip. Interface *C* is formed by the back-to-back assembly of two LRRs between helices of the fourth and fifth repeat. The 2-fold axial symmetry centres between the opposing side chains of H784, which is the first residue of the fourth LRR helix. Most prominent in this interface are phenylalanines 788 and 813, staggered on the fourth and fifth repeat helix, respectively, forming hydrophobic interactions. Associative electrostatic interactions between D789 and R816’ as well as R759 and D846’ additionally stabilize the interface by salt bridges. The buried surface areas of the three interfaces are ∼1650, 350, and 1000 Å^2^, respectively, counting both molecules. As interfaces *A* and *B* complement each other on stabilizing a homodimer formed through intertwined LRR domains, whereas interface *C* mediates the assembly into the decamer, the overall assembly is considered a pentamer of dimers rather than a dimer of pentamers. Notably, no intermolecular interactions within adjacent NACHT domains are seen that would additionally stabilize the quaternary structure.

**Fig. 3.**
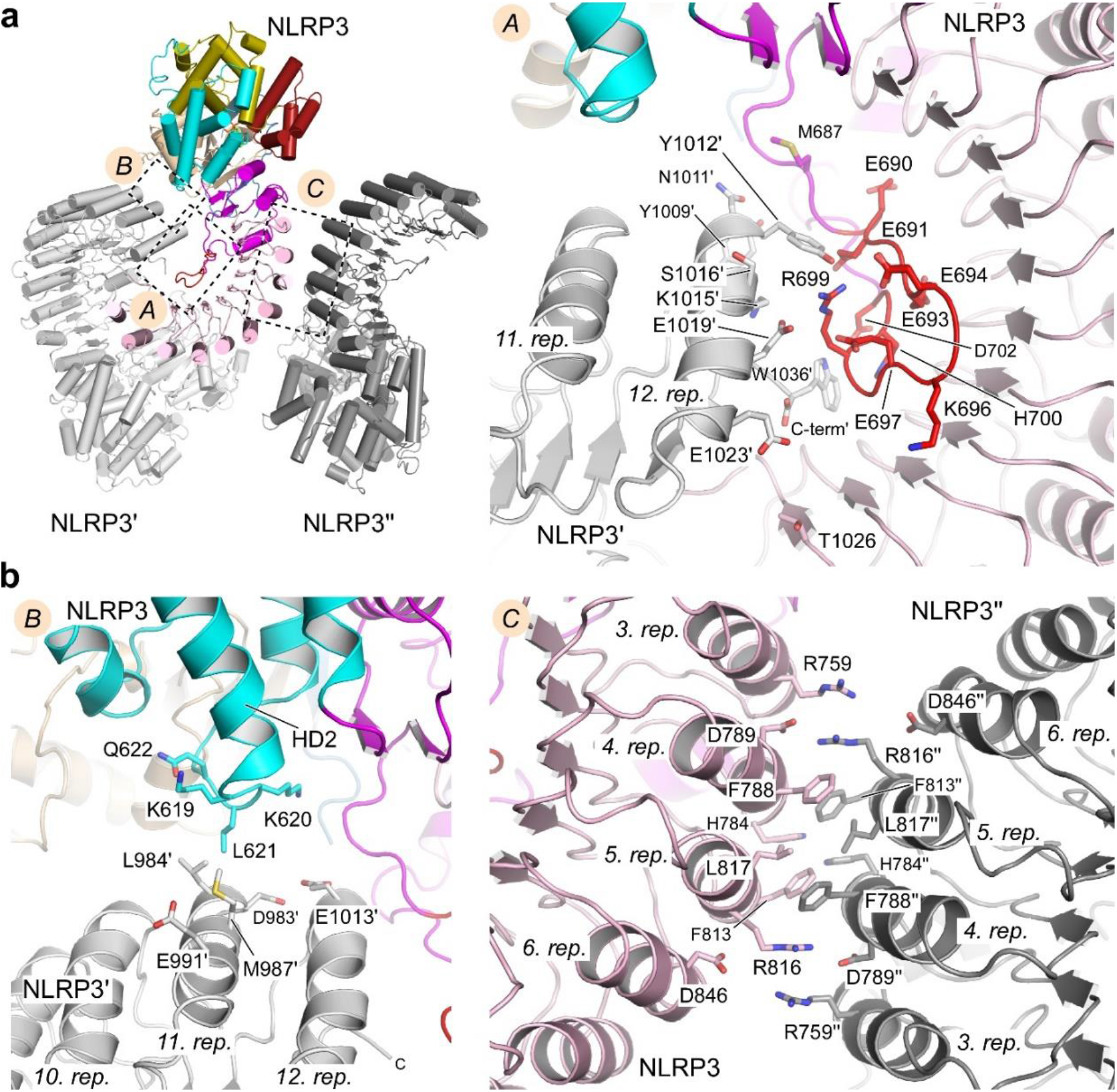
Interfaces in the NLRP3 decamer assembly. **a**, The three unique interfaces in the NLRP3 decamer. Interfaces *A* and *B* form a NLRP3 homodimer with intertwined LRRs, whereas interface *C* is the back-to-back assembly of the LRR leading to the pentamer formation of dimers. **b**, Close up of the individual interfaces derived from density-based modelling. Interacting residues are indicated and coloured according to the subdomain.

## CRID3 binding to NLRP3

The CRID3 inhibitor with a mass of 402 Da binds to a cleft in NLRP3 spanned by subdomains NBD, HD1, WHD, HD2 and trLRR. Surface plasmon resonance showed a dissociation constant of 20 ± 4 nM for CRID3 binding to NLRP3 (Fig. 4a). At the present resolution, the individual parts of the drug can be unambiguously assigned to the Coulomb potential map (Fig. 4b). Still, exact geometries of hydrogen bonds or salt bridges cannot be defined. The central sulfonyl-urea group is located on the backside of the Walker A motif close to the side chains of the two alanines within the G_226_AAGIGKT sequence (Fig. 4c). It is sandwiched between arginines R351 and R578, reaching out from opposing sites from the NBD and HD2 domains. R351 is of particular interest, as it is located at the tip of the central β-strand within the pentamerous parallel β-sheet. This residue is known to sense the phosphorylation state of the bound nucleotide in AAA+ ATPases^22^, transforming a triphosphate-loaded state into a conformational change between subdomains NBD and HD2. The hydrophobic, butterfly-shaped hexahydro-*s*-indacen-4-amine contacts HD1 with its aniline moiety at residues M408 and V414, flanked on one side by T439, Y443 and T524 of WHD and F575 of HD2, and on the other side by M661 of the trLRR. The furan head group at the other end of the drug gets close to P352 of NBD, with its tertiary alcohol group contacting Y632 and L628 of HD2. This terminal group reaches out of the binding crevice whose exit funnel is formed by E369, Q225, G226, P352, V353 (all NBD), L628, Q627 (both HD2), and D662, S658 (both trLRR) (Extended Data Fig. 10). In summary, it is remarkable how CRID3 interacts with residues from five different subdomains, stabilizing their conformational arrangement.

**Fig. 4.**
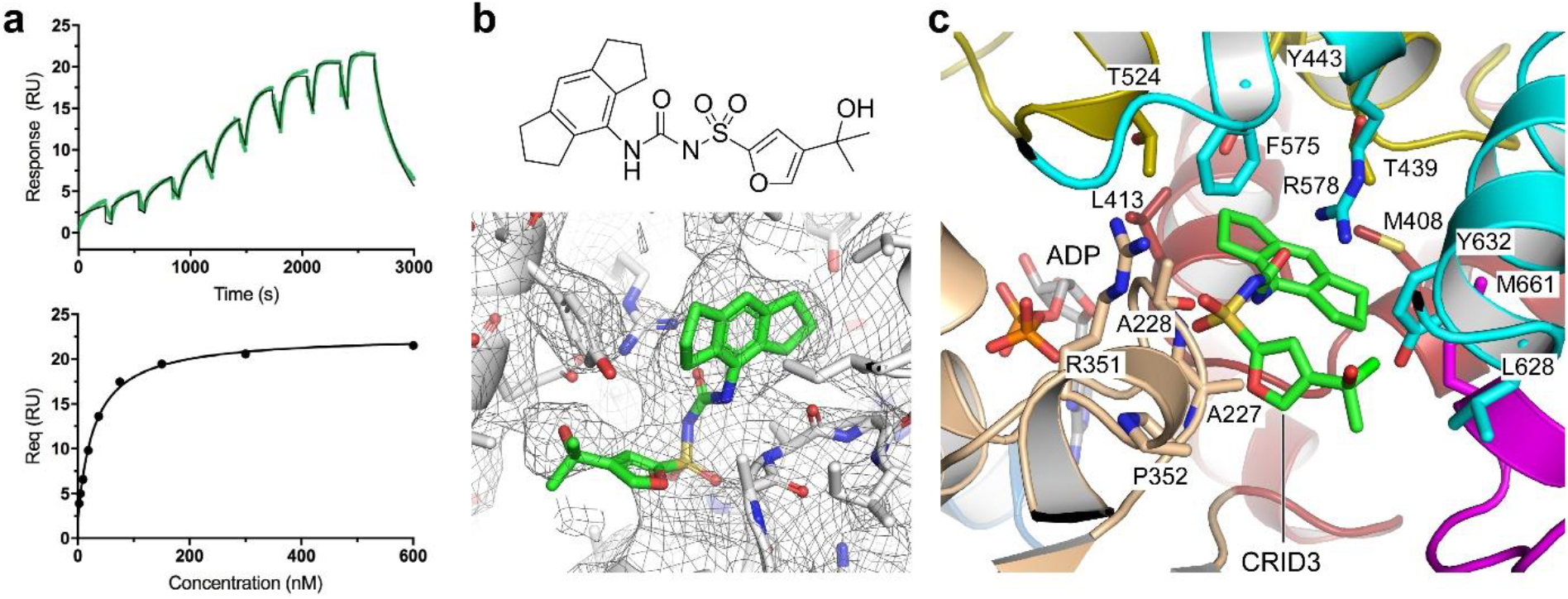
CRID3 binds into a crevice in NLRP3 interacting with five subdomains. **a**, Surface plasmon resonance of the interaction of CRID3 with NLRP3 reveals a *K*_D_ of 19.6 ± 4 nM. **b**, Structure of CRID3 and multibody density map displayed at a 5.0 root mean square deviation (r.m.s.d.) threshold. **c**, Molecular interactions of NLRP3 with CRID3. Residues from five subdomains directly contact CRID3. The central sulfonylurea group is sandwiched between A228, R351 and R578.

## Conclusion

Our structural analysis reveals that full-length, ADP-bound NLRP3 is a decamer. This observation agrees with previous data, showing that before stimulation, NLRP3 proteins form a preassembled complex larger than a megadalton in size^4,23,24^. The decamer appears as a pentamer of dimers, with the C-terminus of NLRP3 entirely buried by the interlocking LRRs. The binding of NEK7 into the concave site of the LRR could be a licensing step for NLRP3 disassembly, promoting its subsequent activation. Five NACHT domains join each side of the sphere, and the PYDs are not resolved in the density map, leaving open whether one or more PYDs are hidden in the decamer cavity. Nevertheless, five PYDs in close proximity are not sufficient to form a stable seed for filament formation^2^. This might be considered a safeguard mechanism against accidental activation but also opens the possibility for a fast-track activation, as only one additional subunit could be sufficient for NLRP3 activation. Interestingly, the activation loops with their two phosphorylation sites S198 and S201 are hidden in the inner side of the decamer, providing yet another protection mechanism.

Our structure of NLRP3 bound to the inhibitor CRID3 resolves a long-lasting question on the binding site and mode of action of this inflammation antagonist. CRID3 ties five subunits of NLRP3 together, presumably precluding the conformational change between NBD/HD1 and WHD/HD2 characteristic for AAA+ activation and ATP binding^15^. Therapeutic intervention to inhibit NLRP3 inflammasome activation could have clinical benefits for conditions where chronic, low-grade inflammation is implicated^11^. Recently, CRID3 derivates reached clinical trials that may in the future find application in rheumatoid- and osteoarthritis, Crohn’s disease, gout, coronary artery diseases, hepatic and pulmonary fibrosis, and non-alcoholic steatohepatitis^25^. NLRP3 inhibitors are first-in-class drugs in inflammatory disease treatment. Knowing their binding site and understanding their mode of action opens a wide field for optimization and the development of other NLR-specific inhibitors as new medicines.

## Acknowledgments

This work was supported by iNEXT-Discovery grant PID 11967 for the NLRP3–CRID3 sample and INSTRUCT grant PID 11340 for the NLRP3 apo sample. We thank Wim Hagen and Felix Weis at EMBL Heidelberg and Alexandre Durand at IGBMC Strasbourg for excellent cryo-EM data recordings. We thank Holger Stark, Göttingen, for initial advice on EM sample preparation. CE thanks Bettina Böttcher for facility access, Ralph Witzgall for JEM2100F access, Gunter Meister for shared infrastructure, Gerhard Lehmann and Norbert Eichner for IT support, and all members of the Structural Biochemistry Group of the Regensburg Center for Biochemistry for help and support. CE and MP acknowledge funding by the Emmy Noether Programme (DFG grant no. EN 1204/1-1 to CE) and SFB 960 (TP-A8 to CE). M.G. and E.L. are funded by the Deutsche Forschungsgemeinschaft under Germany’s Excellence Strategy – EXC2151 – 390873048.

## Author contributions

I.V.H. performed all experiments and data analysis, except for the following: J.M. characterized the oligomeric state in cells. M.M. performed CRID3 binding experiments. R.B. identified Peak 2. I.V.H. and M.P. prepared cryo-EM samples. I.V.H., M.P., G.H. and C.E. processed cryo-EM data. G.H., M.G., and M.P., C.E. built the model. I.V.H., M.P., G.H., C.E. and M.G. interpreted the data. E.L. provided reagents for cell-based assays and valuable discussions. C.E. supervised cryo-EM sample preparation, optimization and data interpretation and contributed to funding. M.G. designed and supervised research and wrote the manuscript. All authors discussed and commented on the manuscript.

## Methods

### Data reporting

No statistical methods were used to predetermine sample size. The experiments were not randomized and the investigators were not blinded to allocation during experiments.

### Cloning, protein expression, and purification

Human NLRP3 (aa 3-1036, UniProt accession code Q96P20), codon-optimized for *Spodoptera frugiperda* was subcloned into an in-house modified pACEBac1 vector containing an N-terminal MBP-tag, followed by a Tobacco etch virus (TEV) protease cleavage site. Recombinant human MBP-NLRP3 was expressed in *Sf9* insect cells, using the Bac-to-Bac system to generate baculovirus infected insect cells. For recombinant protein expression, 1 L of insect cells were supplemented with 30 ml of a viral stock of the second passage. The expression culture was incubated for 72 h at 27°C and 80 rpm, and was subsequently harvested by centrifugation at 2000 rpm for 20 min. The cell pellet was washed with PBS and cells were again spun down at 2000 rpm for 20 min. The resulting cell pellet was either used for subsequent protein purification or frozen in liquid nitrogen and stored at -80°C till further use. For protein purification, a 1 L cell pellet was resuspended in lysis buffer (50 mM Hepes pH 7.5, 150 mM NaCl, 0.5 mM TCEP, 10 mM MgCl_2_, 1 mM ADP) supplemented with 100 µM PMSF, followed by sonication (5 sec on; 10 sec off for 4 min at 40% intensity on ice) and subsequent centrifugation (1 h, 25,000 rpm). The supernatant was filtered through a 0.45 µm syringe filter and applied onto a 5 ml MBPtrap column (GE Healthcare). The column was washed with 10 CV of lysis buffer and the protein was eluted with 5 CV of lysis buffer supplemented with 15 mM maltose. The recombinant protein was digested with TEV-protease overnight at 4°C (ratio TEV to protein of 1:50) and subsequently cross-linked with 0.5 mM bis(sulfosuccinimidyl)-suberate (BS3) for 30 min at 30°C. The reaction was quenched by addition of 100 mM ammoniumhydrogencarbonate for 15 min at 30°C. The cross-linked protein was concentrated to 1/3 of the original volume using a 15 ml spin concentrator (Amicon, 100 kDa cut-off). The cross-linked recombinant protein was further purified by size exclusion chromatography (SEC) on a Superose 6 (S6) increase 3.2/300 column. For purification of the CRID3 bound protein, CRID3 was supplied to the expression culture to final concentration of 0.01 mM, as well as to all buffers. Purified proteins were analysed by SDS-PAGE and negative stain EM.

### Cryo-EM grid preparation, data collection, and processing

The purified and cross-linked recombinant NLRP3 protein was loaded onto glow-discharged EM-grids (Quantifoil, R2/1 Cu-covered with homemade ultrathin carbon support film), incubated for 30 sec and blotted for 5 sec, force 9, at 100% humidity and 4°C prior to plunge freezing into liquid ethane using a Vitrobot mark IV plunge freezer (ThermoFisher). For data set acquisition of apo NLRP3, 9,186 molecular movies were acquired at a cs corrected Titan Krios equipped with a K3 camera, with a total dose of 40.63 e^-^/A^2^ fractionated over 40 frames with a defocus range of 1.5 to 3 µm. For NLRP3–CRID3 data set acquisition, movie frames were collected in counting mode at 0.645 Å per pixel on a Titan Krios equipped with a Quantum K3 camera (Gatan). 14,336 movie frames with a total dose of 45 e^-^/A^2^ over 40 frames were collected at a defocus range of 0.75 to 1.5 µm.

Cryo-EM data processing was done in RELION3.1^26^. Movie frames were aligned using RELION’s own implementation of the MOTIONCORR-2 algorithm. Motion-corrected but non-dose-weighted micrographs were used to determine CTF parameters using CTFFIND 4.1. Initially, 1,588,061 particles were auto-picked in RELION3.1 (Laplacian picker) and subjected to reference-free 2D classification, which yielded 24 2D classes showing high-resolution features accounting for a total of 1,291,183 particles. From these particles an initial model was calculated that was used as reference map for subsequent 3D classifications sorting for globular shaped particles. One selected 3D class showing a well-defined convex LRR interface and accounting for 161,351 particles was further 3D refined, resulting in a reconstruction at 3.9 Å overall resolution (0.143 FSC) for the NLRP3 decamer. A second processing tree that used masked 3D classification and focussed auto-refinement yielded a NLRP3 NACHT domain reconstruction at an overall resolution of 4.1 Å (0.143 FSC). A third reconstruction of a NLRP3 monomer at an overall resolution of 4.8 Å (0.143 FSC) was generated through a multibody refinement approach.

### Model building and refinement

For model building, four different density maps were generated by focused classifications, masked refinements and multibody refinements that were optimized for the NACHT domain, the LRR domain, the acidic loop, and the equatorial LRR back-to-back interface, respectively. All maps were generated using RELION3.1^26^. The bulk of the model building was performed using a multibody refinement map comprising one monomer of NLRP3 and details were adjusted by overlaying the focused maps. The NACHT domain was separated into its five subdomains FISNA, NBD, HD1, WHD, and HD2 based on the NLRP3–NEK7 complex structure (6NPY)^15^. Each subdomain was stripped for the α-helical and β-strand secondary structure elements only, while loops and flexible regions were omitted. A homology model of the canonical part of the LRR (initially comprising residues 720-1036; later adjusted to 743-1036) was generated by SWISS-Model^27^ using the high-resolution structures of the porcine ribonuclease inhibitor protein (2BNH)^16^.

The initial model was built by placing the stripped subdomains NBD, HD1, WHD and cnLRR into the density using rigid-body fitting in Chimera X^28^. Divergent parts of these subdomains were manually adjusted in Coot^29^ and the loop sections were built to fit the density. The map produced by focused refinement of a single NACHT domain was used for this step. As the templates for FISNA, HD2 and the transition region connecting to the cnLRR did not fit sufficiently well, homology modelling using SWISS-Model based on the crystal structures of NOD2 (5IRN)^17^ and NLRC4 (KXF)^18^ was performed. The helices and β-strands of all subunits were placed and built with confidence in their register using bulky residues (e.g., R141, F148; Y572, F575, R578, W611, Y632; F650, F729) as sequence markers correlated with secondary structure predictions. The extra density of the CRID3 and ADP ligands was clearly visible and allowed us to confidently place these ligands into the model (Extended Data Fig. 5).

Already in the first reconstructions, we identified an elongated stretch of extra density at the concave interface *A* that had features similar to a polypeptide chain. A focused refinement of this region revealed that the density is caused by the NLRP3-specific acidic loop starting from F683 to H724, which was manually built de novo using Coot. The resulting model of NLRP3 bound to CRID3 and ADP was subjected to multiple rounds of manual adjustment and real-space refinement in Coot, PHENIX^30^ and ISOLDE^31^. The interfaces were refined using a NLRP3 trimer. The central monomer of this trimer was then assembled into a decamer structure by fitting 10 monomers into the highest resolution decamer map. The final model showed reasonable stereochemistry as assessed by MolProbity^32^ (Extended Data Table 1). Interaction analysis of the decamer interfaces was performed visually and using PDBePISA^33^. Figures representing the structures and maps were prepared using PyMOL and UCSF Chimera X.

### Multi-angle light scattering

For molecular-mass determination by multi-angle light scattering (MALS), protein samples were injected into a Superose 6 (10/300 GL) gel-filtration column equilibrated with gel filtration buffer. The chromatography system was attached to a three-angle light scattering detector (miniDAWN, Wyatt) and a refractive index detector (Optilab T-rEX, Wyatt). Data were collected every 0.5 s with a flow rate of 0.5 ml/min. Data analysis was carried out using the ASTRA V software.

### Surface plasmon resonance spectroscopy

Surface plasmon resonance (SPR) experiments were performed on a Biacore 8K (GE Healthcare) device. The system was flushed with running buffer (10 mM HEPES pH 7.4, 200 mM NaCl, 0.5 mM ADP, 0.5 mM tris(2-carboxyethyl)phosphine (TCEP), 2 mM MgCl_2_, 1 g/L carboxymethyl dextran hydrogel (CMD), 0.05% Tween20, 2% DMSO) at 25°C. A streptavidin functionalized sensor chip (Series S Sensor Chip SA, Cytiva) was conditioned with three consecutive injections of 1 M NaCl in 50 mM NaOH (10 μL/min) for 1 min as described^34^. Biotinylated NLRP3 NACHT-trLRR protein from HEK cell expression was immobilized at 2 μL/min for 900 s. The flow system was washed using 50% isopropanol in 1 M NaCl and 50 mM NaOH. Free streptavidin binding sites were blocked by four consecutive injections of Biotin-PEG (1000 nM, M_n_ 2,300 Da) for 2 min at 10 µL/min. For kinetic binding measurements in the single cycle mode, increasing concentrations of 5 to 600 nM CRID3 were injected at 30 μL/min (association 240 s, dissociation 60/360 s). Data were collected at a rate of 10 Hz. The binding data were double referenced by blank cycle and reference flow cell subtraction. Data were corrected by a 4-point solvent correction. For kinetic binding experiments, processed data were fitted to a 1:1 interaction model using the Biacore Insight Evaluation Software (version 3.0.12.15655).

### Data availability

The cryo-EM density reconstructions and models were deposited with the Electron Microscopy Data Bank (EMDB) (accession codes EMD-xxxxx for the NLRP3 decamer and EMD-xxxxy for the NLRP3:ADP:CRID3 decamer) and with the Protein Data Bank (PDB) (accession code XXXX). All data are available in the Article or its supplementary files. Source data are provided with this paper.

## Extended Data Figures

**Extended Data Fig. 1.**
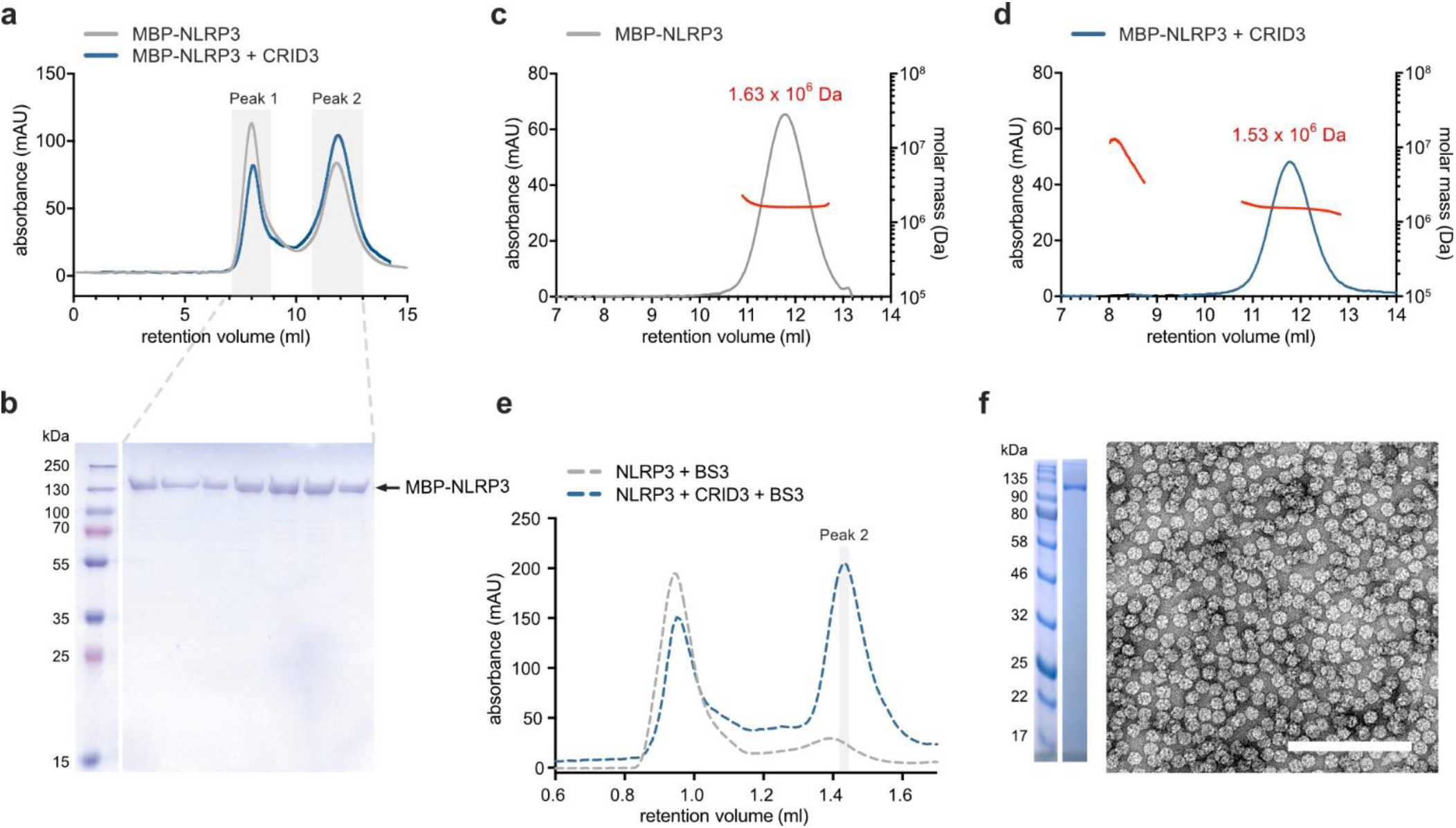
Preparation of full length, wild type, human NLRP3 for EM analyses. **a**, Analytical size-exclusion chromatography of recombinant, human MBP-NLRP3 (3-1036) reveals two elution peaks, one close to the void volume (Peak 1) and one at a size larger than 1 MDa (Peak 2). The peak ratio is shifted from Peak 1 under standard buffer conditions (50 mM HEPES pH 7.5, 150 mM NaCl, 10 mM MgCl_2_, 1 mM ADP, 0.5 mM TCEP) towards Peak 2 by addition of CRID3 (10 µM CRID3). Analytical SEC runs were performed on a Superose 6 Increase 10/300 GL column with 700 µl applied at a concentration of 3 mg/ml. **b**, Coomassie-stained SDS PAGE analysis of purified MBP-NLRP3. **c**, SEC-MALS experiment of MBP-NLRP3 Peak 2 plus ADP on a Superose 6 Increase 10/300 GL column. The calculated mass of an MBP-NLRP3 monomer is 158 kDa. **d**, SEC-MALS experiment of MBP-NLRP3 Peak 2 plus ADP plus CRID3 on a Superose 6 Increase 10/300 GL column. The average apparent mass of Peak 2 from both experiments equals 10-times the calculated mass of the protein. **e**, Analytical SEC of recombinant, human NLRP3 (3-1036) after TEV digestion using a Superose 6 Increase 3.2/300 column. **f**, Negative-stain micrograph of TEV-digested, gel filtered NLRP3 plus CRID3 plus BS3 Peak 2 elution fraction that was used for subsequent cryo-EM grid preparation. An SDS-PAGE analysis of the protein w/o BS3 is shown left. The scale-bar equals 200 nm.

**Extended Data Fig. 2.**
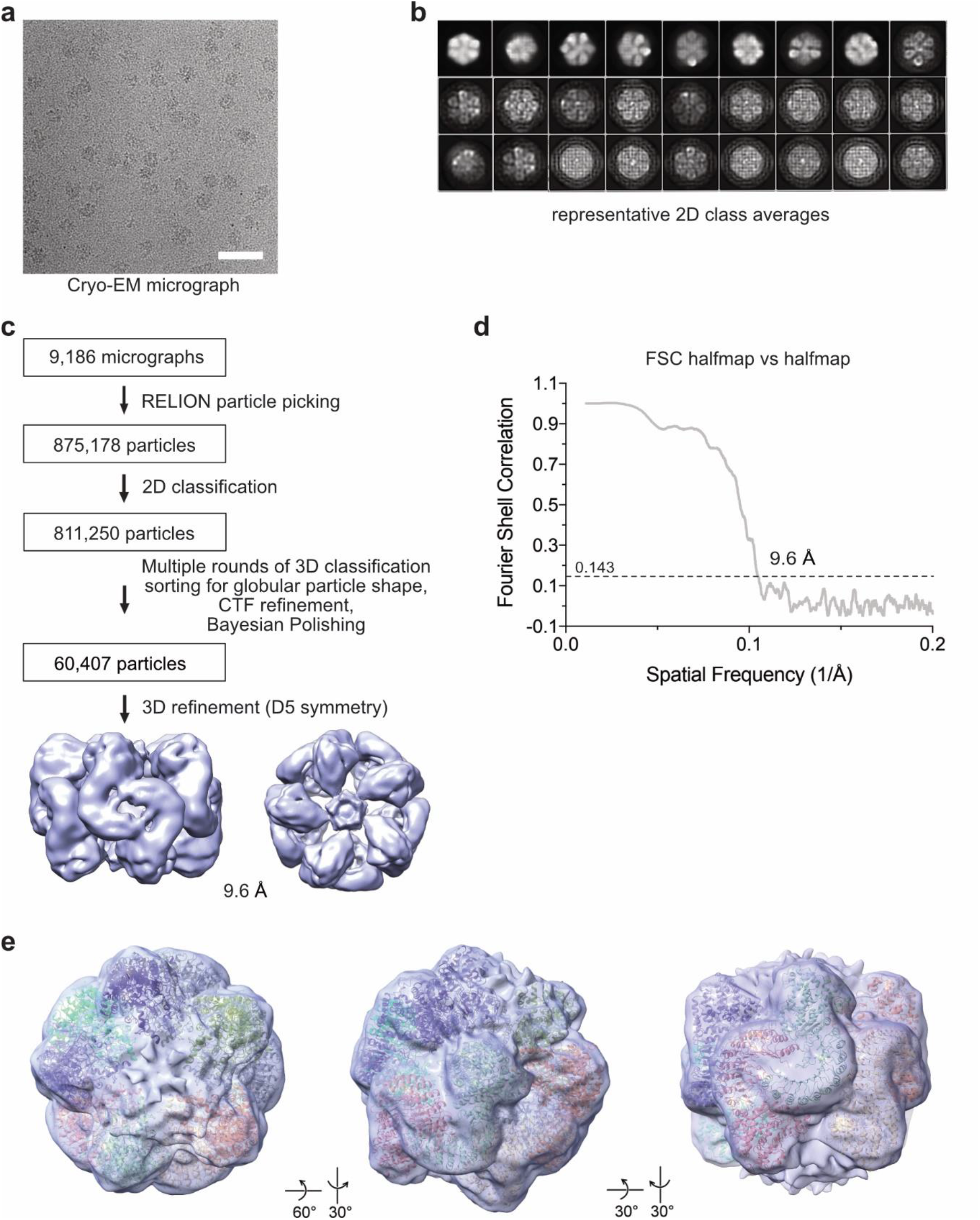
Cryo-EM data processing of NLRP3. **a**, Representative cryo-EM micrograph of the human NLRP3 oligomer eluting as Peak 2 (replicated more than 35,000 times). Scale bar, 50 nm. **b**, Representative 2D class averages. **c**, Processing tree describing particle classification of the NLRP3 cryo-EM data in RELION. **d**, Fourier Shell Correlation (FSC) of the NLRP3 decamer particles. **e**, Top, tilted, and side views of the cryo-EM density of the NLRP3 protein w/o CRID3 at ∼10 Å resolution. The 3.9 Å structure of the NLRP3–CRID3 complex is fitted into the density map showing that the apo and the inhibitor bound NLRP3 decamer structures do not exhibit large conformational rearrangements.

**Extended Data Fig. 3.**
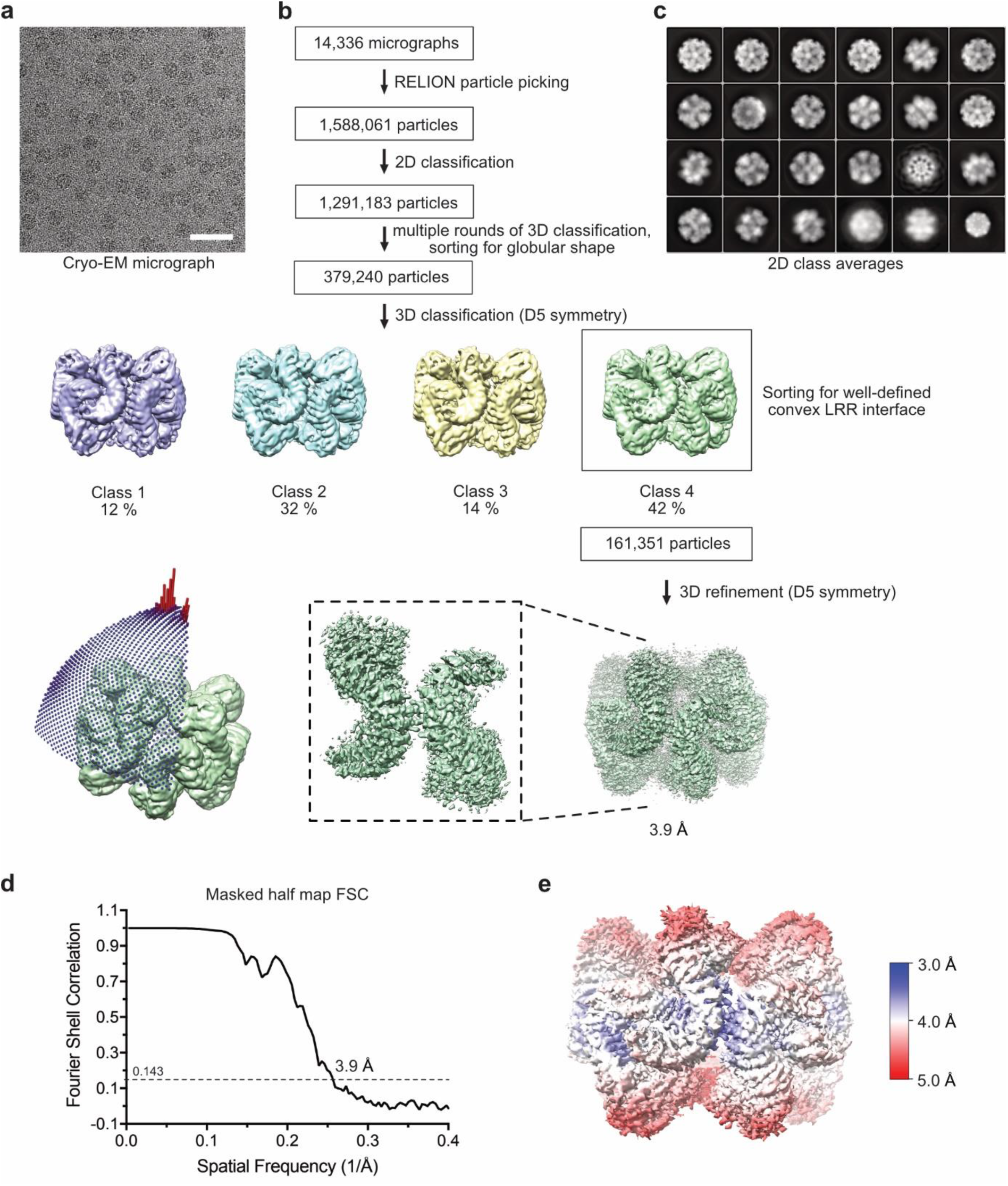
Cryo-EM data processing of the NLRP3–CRID3 complex. **a**, Representative cryo-EM micrograph of the human NLRP3 decamer in complex with CRID3 (replicated more than 20,000 times). Scale bar, 50 nm. **b**, Processing tree of cryo-EM data as described in the Methods section. **c**, Representative 2D class averages. **d**, Masked FSC of particles in the decamer reconstruction. **e**, Reconstruction coloured by the local resolution as estimated by RELION3.1^26^.

**Extended Data Fig. 4.**
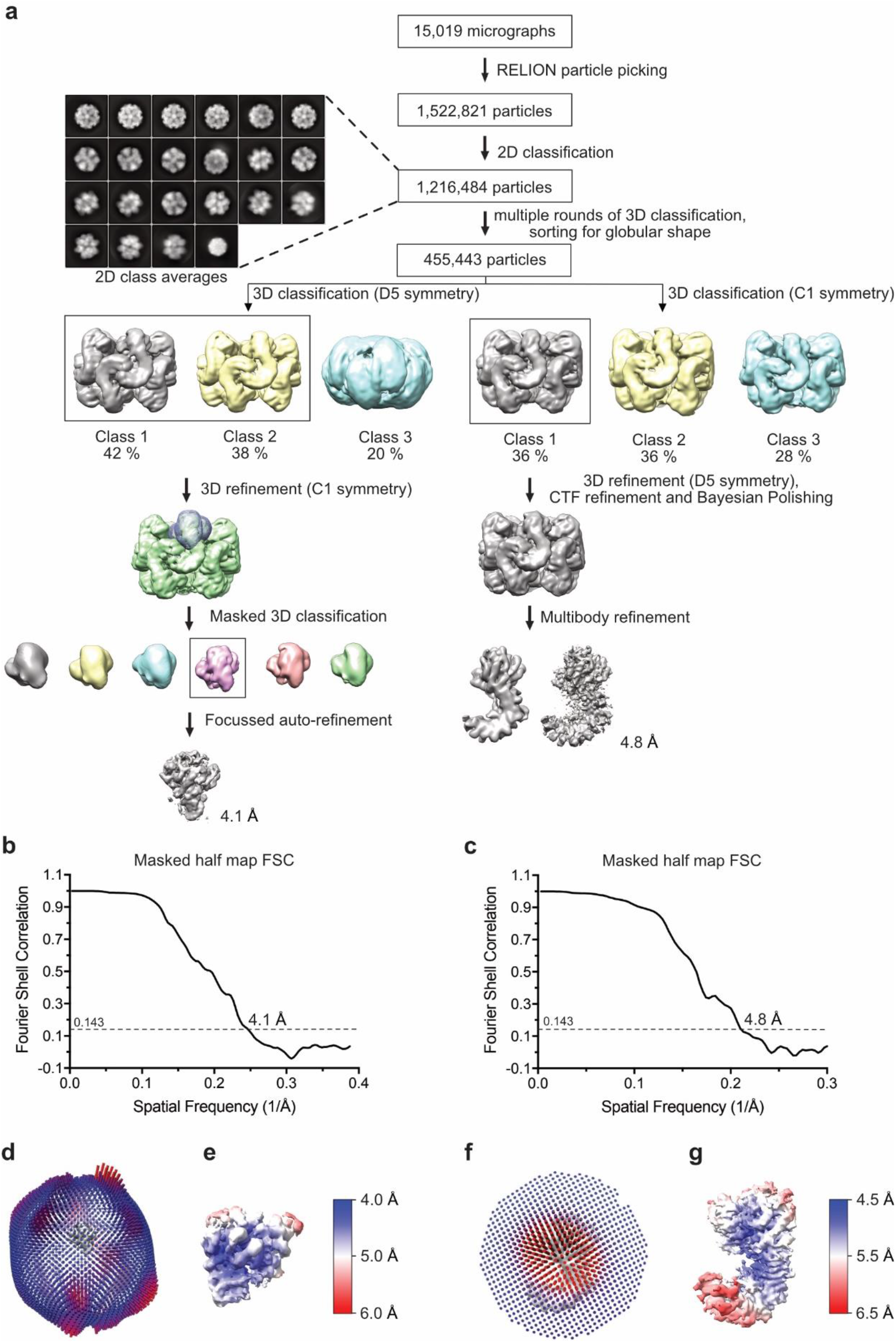
Cryo-EM data processing of the NLRP3–CRID3 complex. **a**, Processing tree of cryo-EM data according to the focussed refinement and multibody refinement workflow. **b, c**, Masked FSC of particles in the focussed (b) and multibody (c) refinement reconstruction. **d**, Orientation distribution of particles in the focussed NACHT reconstruction. **e**, Local resolution estimation of the final focussed NACHT reconstruction. **f**, Orientation distribution of particles in the multibody refinement reconstruction. **g**, Local resolution estimation of the final multibody reconstruction.

**Extended Data Fig. 5.**
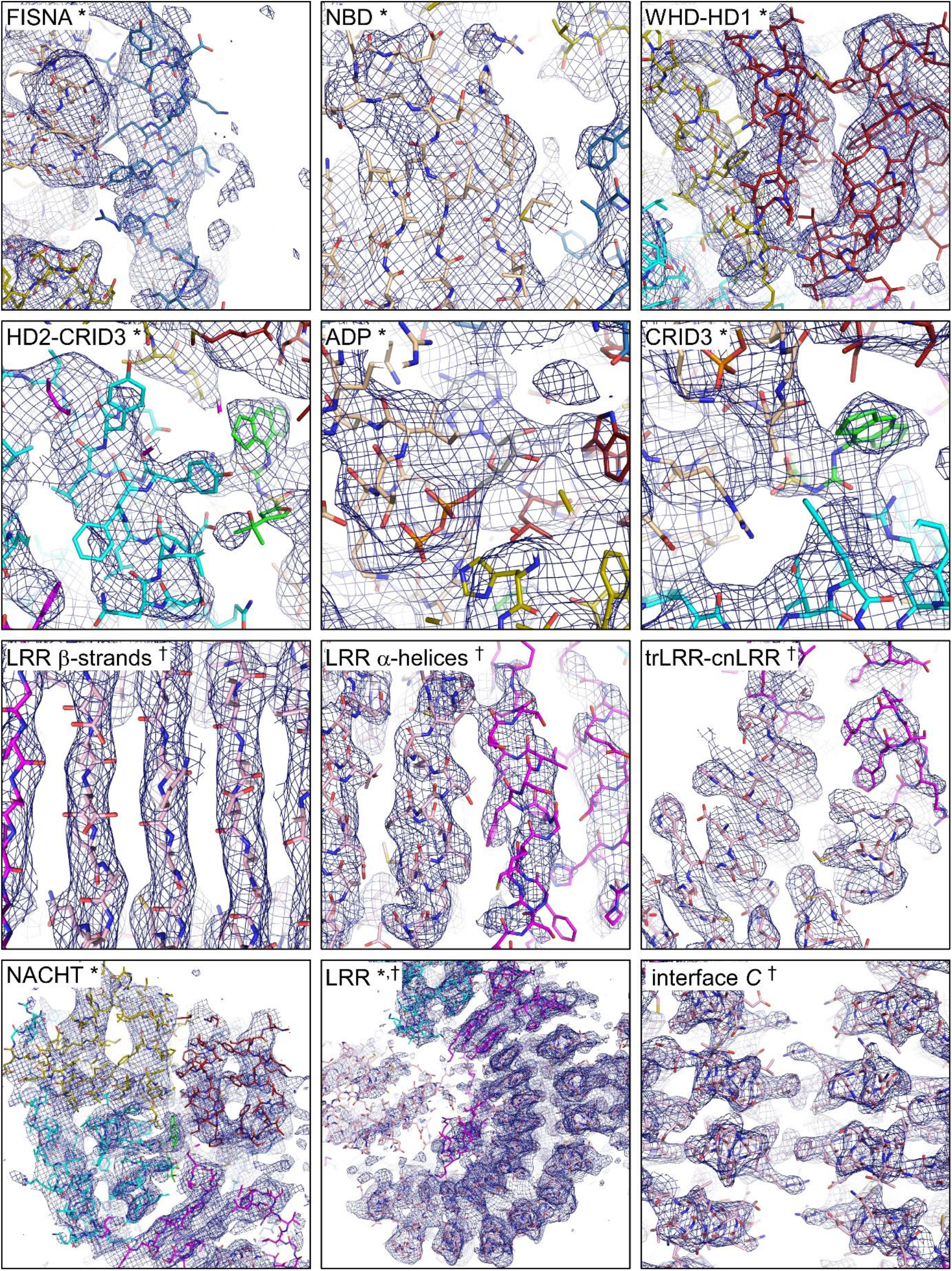
Quality of cryo-EM densities. Sections of focused-refined NLRP3–CRID3 cryo-EM density overlaid with their respective atomic models. Densities are shown as a blue mesh, and sticks are shown for the structure model coloured as in Fig. 2a. The labels ‘*’ and ‘^†^’ refer to the ‘multibody refine monomer’ and ‘best decamer’ density, respectively.

**Extended Data Fig. 6.**
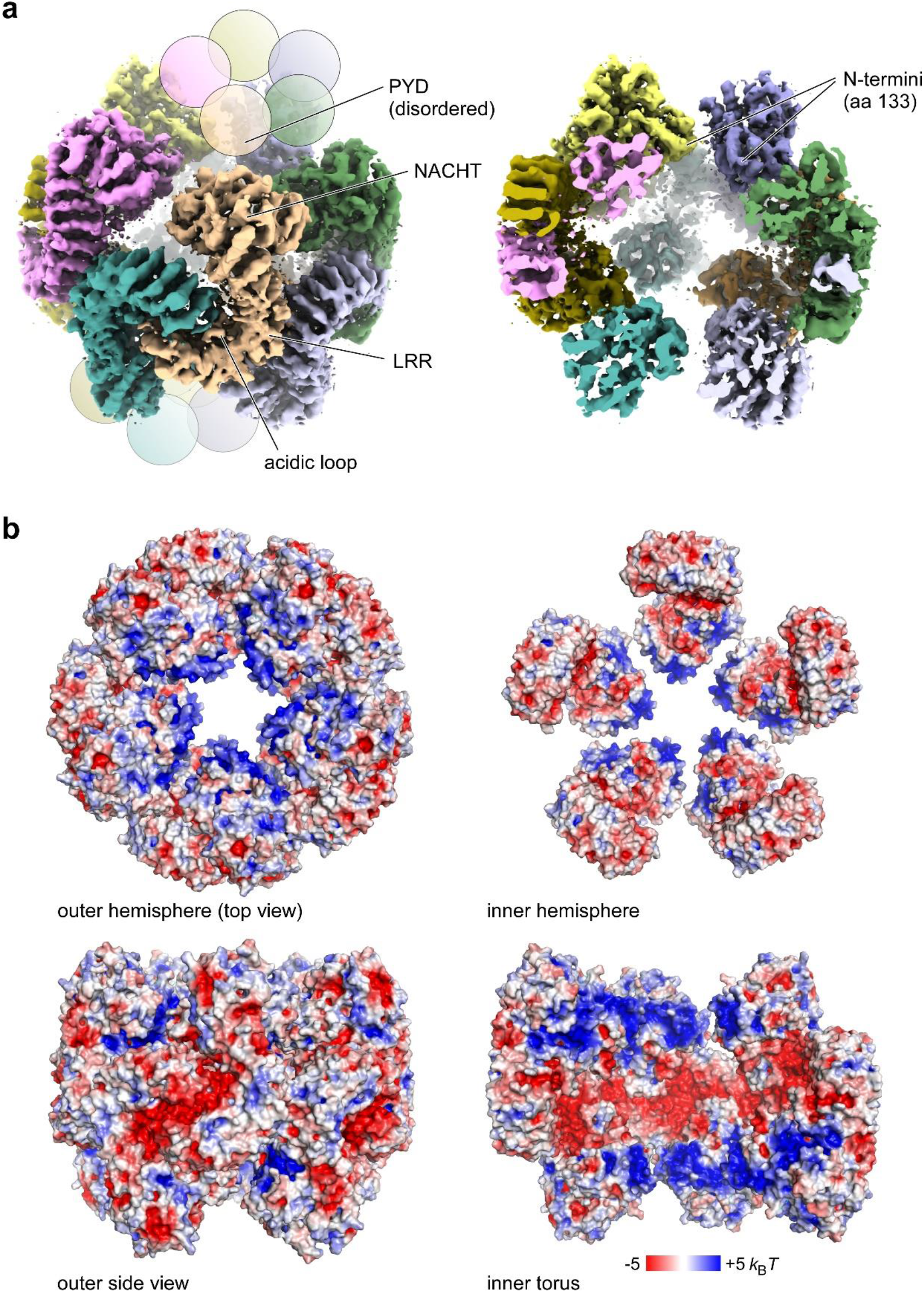
Features of the NLRP3 decamer structure. **a**, Tilted view on the cryo-EM density of the NLRP3–CRID3 decamer structure with the NACHT and LRR domains indicated (left). The acidic loop mediating the intertwined LRR assembly is labelled. The PYDs at the apical site of the sphere are disordered in the cryo-EM map but drawn here as balls. The density map is shown at a threshold level of 0.257. (Right) Cross section of the cryo-EM density shown left. The N-termini of the resolved structure (aa 133) pointing toward the hollow of the sphere are indicated. **b**, Four views on the electrostatic surface potential of the NLRP3–CRID3 decamer structure. Whereas the basic cluster at the N-terminus of the FISNA forms a positively charged rim of the decamer cavity, the inner torus is negatively charged.

**Extended Data Fig. 7.**
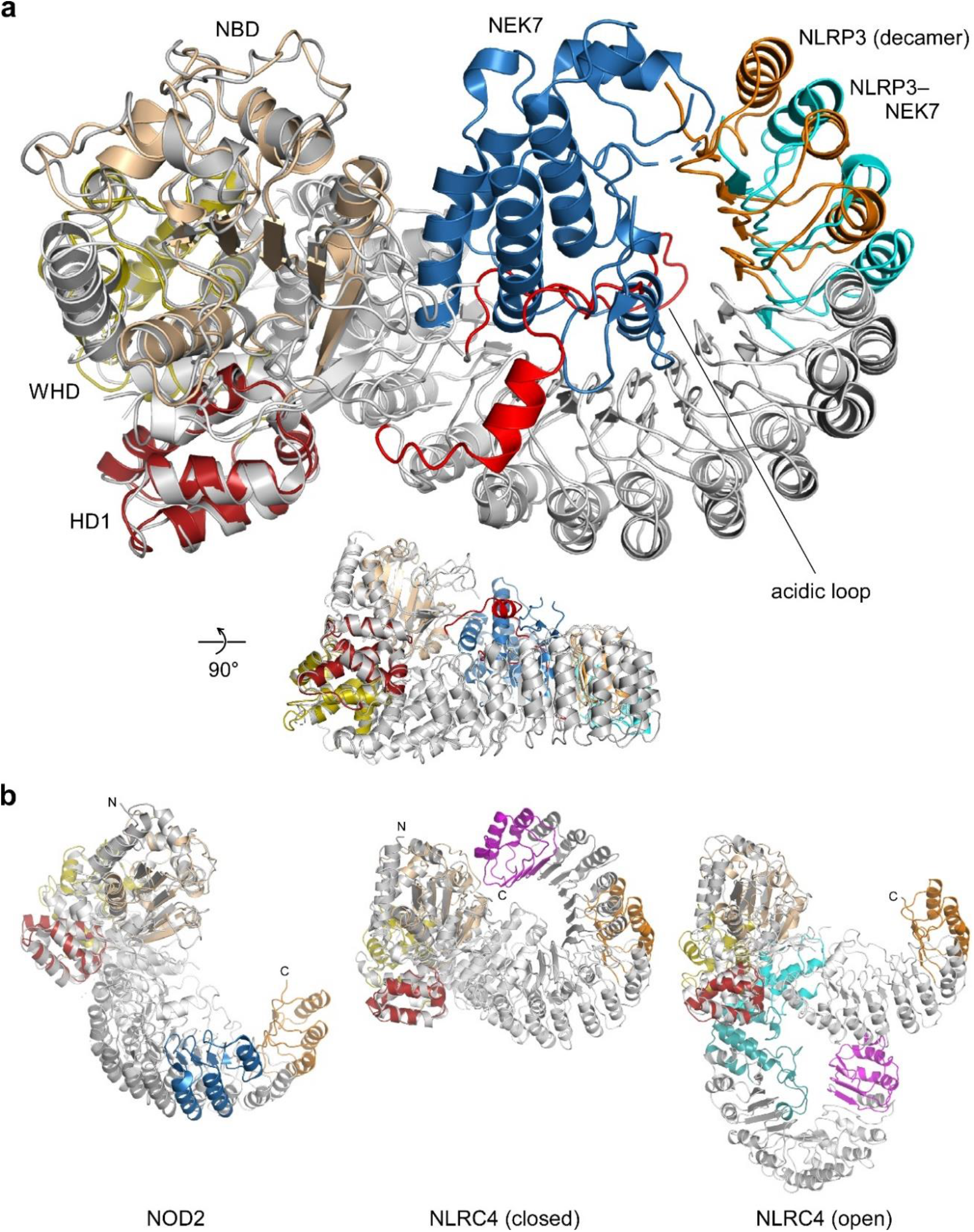
The LRR in the NLRP3 decamer adopts a partially open conformation. **a**, Overlay of NLRP3 from the decamer structure with the NLRP3–NEK7 complex structure (6NPY)^15^. Proteins were aligned for the NBD-HD1 subdomains to visualize the conformation of the LRR relative to the NACHT domain. For better clarity, only the last three repeats of the LRRs are coloured. The C-terminal lobe of the kinase NEK7 (blue) adopts the space in the concave site of the LRR that is occupied in the NLRP3 decamer structure by the acidic loop (red). The LRR in the NLRP3–NEK7 structure is only slightly more open (∼½ repeat) compared to the NLRP3 decamer structure. **b**, Overlay of NLRP3 from the decamer assembly with NLR family proteins NOD2 (5IRN)^17^, monomeric NLRC4 (4KXF)^18^, and the disc-like NLRC4 (3JBL)^35^ structure. Protein structures were aligned as in a. The last three repeats are coloured blue (NOD2) or magenta (NLRC4).

**Extended Data Fig. 8.**
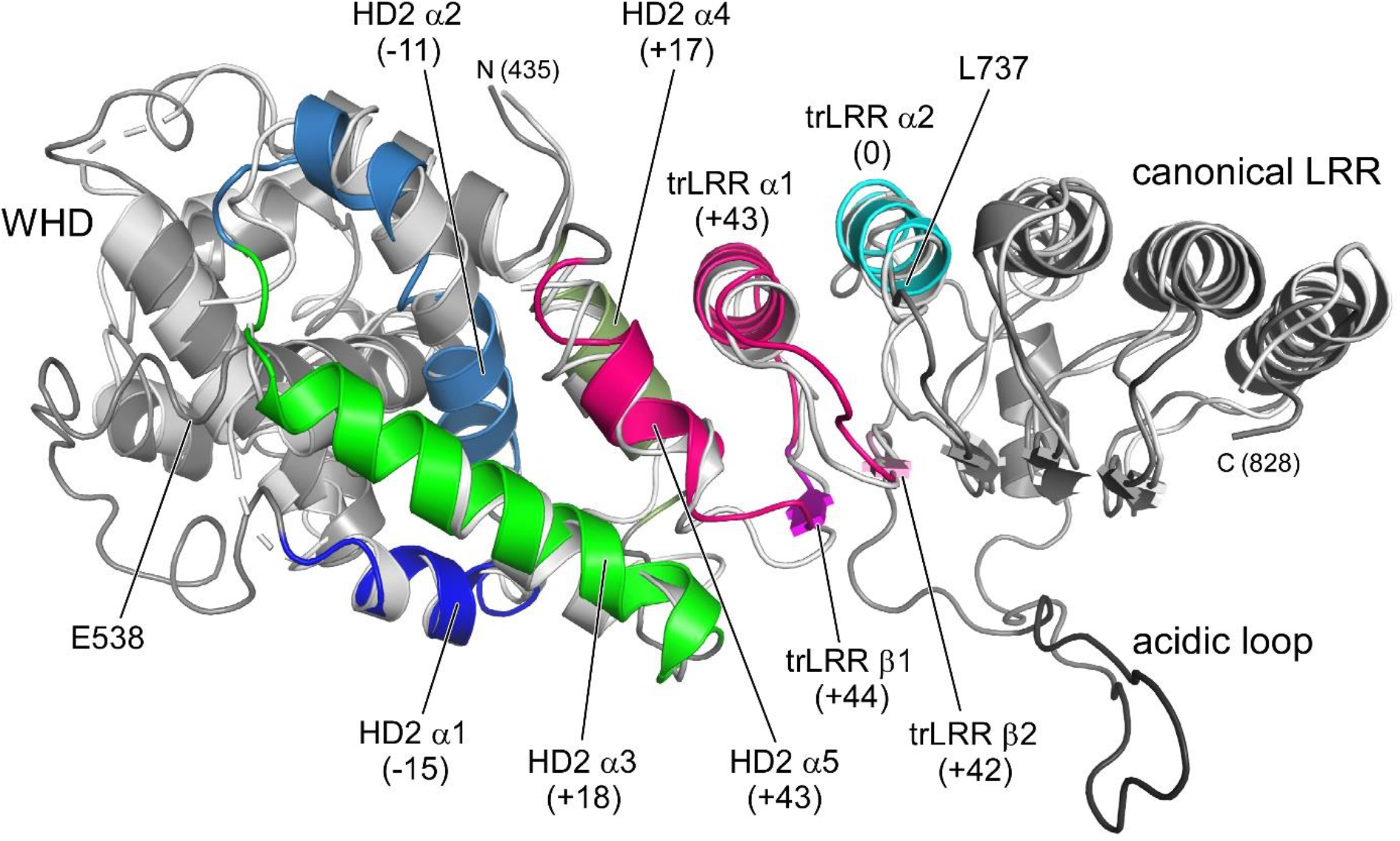
Transitions in HD2 and trLRR between the NLRP3 decamer and the NLRP3–NEK7 structure. Overlay of residues 435-828 between our structure (dark grey/coloured) and the NLRP3–NEK7 structure (6NPY, light grey)^15^ showing WHD, HD1, trLRR and 3 repeats of the canonical LRR. There is a shift in the amino acid register starting from position 538 that varies between -15 residues in the first helix of HD2 and +44 residues in the first β-strand of the trLRR. The register synchronizes again at L737 at the beginning of the canonical LRR domain. Segments are coloured according to the amino acid register shifts.

**Extended Data Fig. 9.**
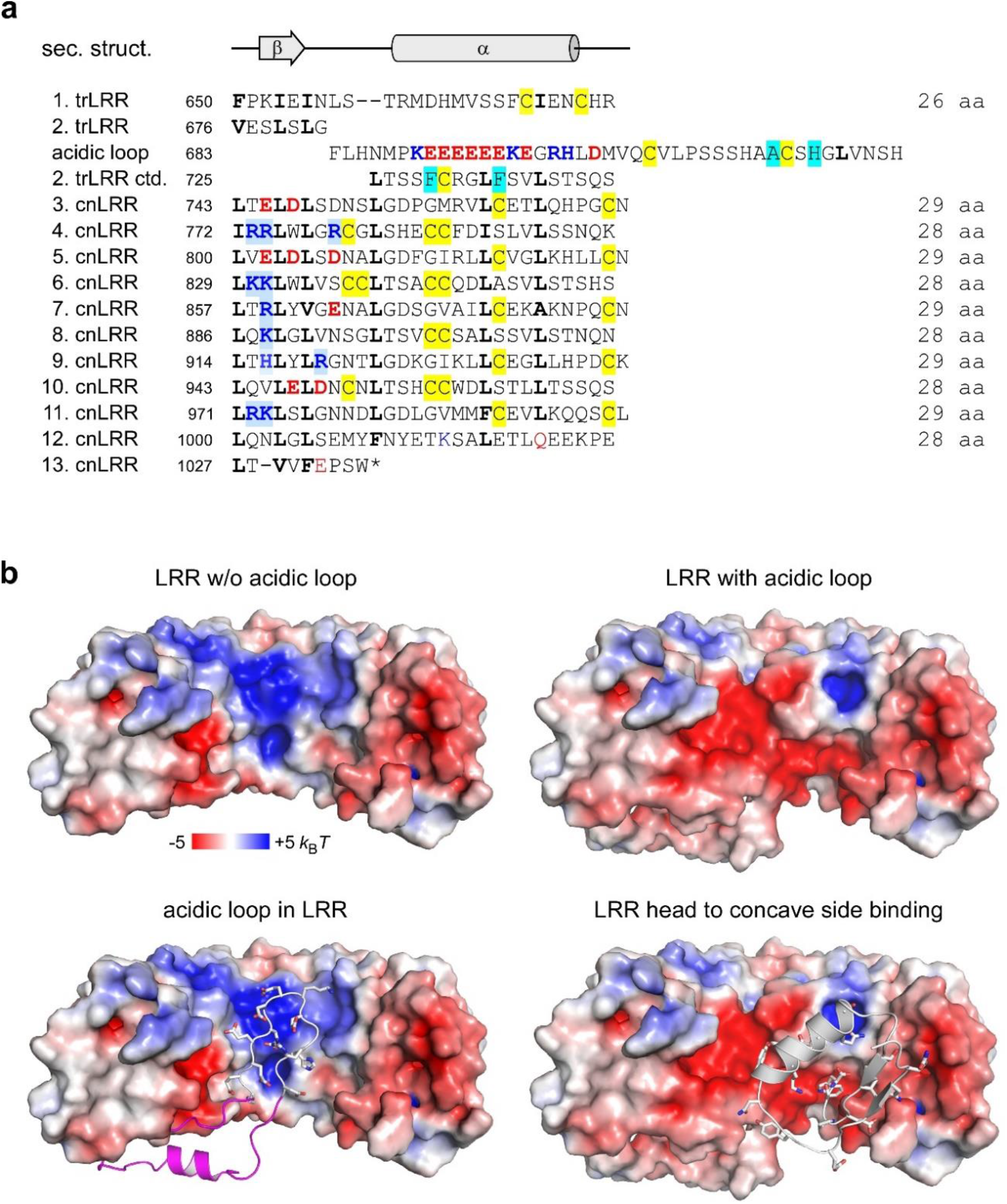
Arrangements of the transition LRR and the canonical LRR. **a**, Sequence alignment of individual repeats of the LRR domain. The transition LRR (trLRR) starts at position F650 with the FxxIxI motif and a 26-aa repeat. An LRR-mismatching region of 42 residues interrupts the conventional fold from residue F683 on, forming a flexible, acidic loop. A highly charged stretch of 14 residues (689-702, theoretical pI 4.4) with acidic residues at the tip binds into the concave side of the LRR. The canonical LRR (cnLRR) starts at position L743 and contains 10 repeats of a prototypic 28/29 residue alteration^35^. Charged residues in the concave surface of the LRR are coloured blue and red. Leucine residues or homologous hydrophobic residues at LRR-defining positions are indicated bold, and cysteines are boxed yellow. Mismatching residues that preclude a canonical LRR fold are boxed cyan. Secondary structure elements of a canonical LRR fold are indicated at the top. **b**, Electrostatics of the LRR–acidic loop–LRR’ interaction. On the left side is the acidic loop interaction in the LRR without the loop (650-1036, Δ683-727) shown. On the right is the interaction of the C-terminal repeat (998’-1036’) of the cognate LRR binding into the concave LRR side shown.

**Extended Data Fig. 10.**
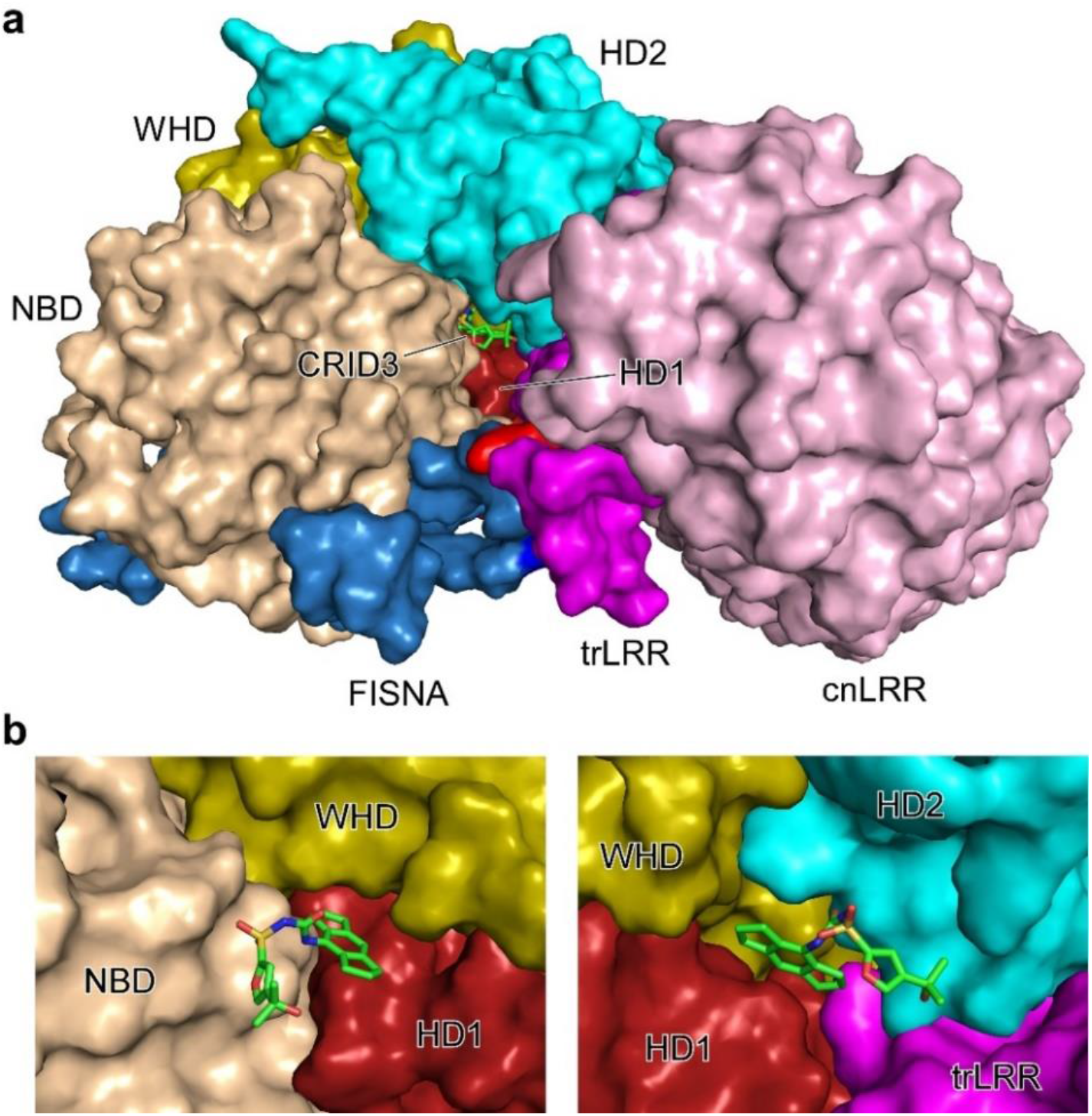
The CRID3-binding site in NLRP3. **a**, CRID3 binds into a deep crevice that is spanned by subdomains NBD, HD1, WHD, HD2 and trLRR. Only the tertiary alcohol group reaches out of this binding cleft. **b**, Close-up of CRID binding to subdomains NBD, HD1 and WHD (left) and HD1, WHD, HD2 and trLRR (right). To visualize the binding sites, the other subdomains in each case were omitted.

**Extended Data Table 1.** Cryo-EM data collection and processing.

| Sample | NLRP3–CRID3 decamer | NLRP3 decamer |
| --- | --- | --- |
| Data acquisition |  |  |
| Voltage | 300 kV | 300 kV |
| Magnification | 130,000 | 81,000 |
| Pixel size | 0.645 Å | 0.862 Å |
| Electron dose | 45 e<sup>-</sup>/Å<sup>2</sup> | 40.63 e<sup>-</sup>/Å<sup>2</sup> |
| Frame number | 40 | 40 |
| Defocus range | 0.75 to 1.5 μm | 1.5 to 3 μm |
| Data Processing |  |  |
| EMDB code | XXXXXX |  |
| PDB code | YYYYY |  |
| Final number of particles | 161,351 |  |
| Map resolution at 0.143 FSC | 3.9 Å |  |
| Local resolution range | 3.9 – 4.8 Å |  |
| Map sharpening B-factor | -150 Å<sup>2</sup> |  |
| Model composition (per monomer of the NLRP3–CRID3 decamer) |  |  |
| Initial PDBs used | 6NPY, 2BNH, 5IRN, 4KXF |  |
| No. of atoms | 7055 |  |
| Protein | 7000 |  |
| Ligands | 27 (ADP)<br>28 (CRID3) |  |
| Mean B factors (Å<sup>2</sup>) |  |  |
| Protein | 117 |  |
| Ligand | 112 |  |
| Model validation |  |  |
| MolProbity score | 2.37 |  |
| Clash score | 17.05 |  |
| Rotamer outliers (%) | 0.13 |  |
| C-beta outliers | 0 |  |
| Ramachandran plot |  |  |
| Favored | 753 |  |
| Outliers | 5 |  |

## References

1. Schroder, K. & Tschopp, J. The inflammasomes. Cell 140, 821–832 (2010).

2. Swanson, K. V., Deng, M. & Ting, J. P. The NLRP3 inflammasome: molecular activation and regulation to therapeutics. Nat. Rev. Immunol 19, 477–489 (2019).

3. Latz, E., Xiao, T. S. & Stutz, A. Activation and regulation of the inflammasomes. Nat. Rev. Immunol. 13, 397–411 (2013).

4. Song, N. et al. NLRP3 phosphorylation is an essential priming event for inflammasome activation. Mol. Cell 68, 185–197 (2017).

5. Schmid-Burgk, J. L. et al. A Genome-wide CRISPR (Clustered Regularly Interspaced Short Palindromic Repeats) Screen Identifies NEK7 as an Essential Component of NLRP3 Inflammasome Activation. J. Biol. Chem. 291, 103–109 (2016).

6. Shi, H. et al. NLRP3 activation and mitosis are mutually exclusive events coordinated by NEK7, a new inflammasome component. Nat. Immunol. 17, 250–258 (2016).

7. He, Y., Zeng, M. Y., Yang, D., Motro, B. & Núñez G. NEK7 is an essential mediator of NLRP3 activation downstream of potassium efflux. Nature 530, 354–357 (2016).

8. Mangan, M. S. J. et al. Targeting the NLRP3 inflammasome in inflammatory diseases. Nat. Rev. Drug Discov. 17, 588–606 (2018).

9. Laliberte, R. E. et al. Glutathione s-transferase omega 1-1 is a target of cytokine release inhibitory drugs and may be responsible for their effect on interleukin-1beta posttranslational processing. J. Biol. Chem. 278, 16567–16578 (2003).

10. Primiano, M. J. et al. Efficacy and pharmacology of the NLRP3 inflammasome inhibitor CP-456,773 (CRID3) in murine models of dermal and pulmonary inflammation. J. Immunol. 197, 2421–2433 (2016).

11. Harrison, D. et al. Discovery of a series of ester-substituted NLRP3 inflammasome inhibitors. Bioorg. Med. Chem. Lett. 30:127560 (2020).

12. Coll, R. C. et al. A small-molecule inhibitor of the NLRP3 inflammasome for the treatment of inflammatory diseases. Nat. Med. 21, 248–255 (2015).

13. Vande Walle, L. et al. MCC950/CRID3 potently targets the NACHT domain of wild-type NLRP3 but not disease-associated mutants for inflammasome inhibition. PLoS Biol. 17:e3000354 (2019).

14. Corcoran, S. E., Halai, R. & Cooper, M. A. Pharmacological inhibition of the Nod-like receptor family Pyrin domain containing 3 inflammasome with MCC950. Pharmacol. Rev. 73, 968–1000 (2021).

15. Sharif, H. et al. Structural mechanism for NEK7-licensed activation of NLRP3 inflammasome. Nature 570, 338–343 (2019).

16. Kobe, B. & Deisenhofer, J. Mechanism of ribonuclease inhibition by ribonuclease inhibitor protein based on the crystal structure of its complex with ribonuclease A. J. Mol. Biol. 264, 1028– 1043 (1996).

17. Maekawa, S., Ohto, U., Shibata, T., Miyake, K. & Shimizu, T. Crystal structure of NOD2 and its implications in human disease. Nat. Commun. 7:11813 (2016).

18. Hu, Z. et al. Crystal structure of NLRC4 reveals its autoinhibition mechanism. Science 341, 172– 175 (2013).

19. Tenthorey, J. L. et al. The structural basis of flagellin detection by NAIP5: A strategy to limit pathogen immune evasion. Science 358, 888–893 (2017).

20. Erzberger, J. P. & Berger, J. M. Evolutionary relationships and structural mechanisms of AAA+ proteins. Annu. Rev. Biophys. Biomol. Struct. 35, 93–114 (2006).

21. Hoss, F. et al. Alternative splicing regulates stochastic NLRP3 activity. Nat. Commun. 10:3238 (2019).

22. Wendler, P., Ciniawsky, S., Kock, M. & Kube, S. Structure and function of the AAA+ nucleotide binding pocket. Biochim. Biophys. Acta 1823, 2–14 (2012).

23. Compan, V. et al. Cell volume regulation modulates NLRP3 inflammasome activation. Immunity 37, 487–500 (2012).

24. Zhang, Z. et al. Protein kinase D at the Golgi controls NLRP3 inflammasome activation. J. Exp. Med. 214, 2671–2693 (2017).

25. Schwaid, A. G. & Spencer, K. B. Strategies for Targeting the NLRP3 Inflammasome in the Clinical and Preclinical Space. J. Med. Chem. 64, 101–122 (2021).

## References

26. Zivanov, J. et al. New tools for automated high-resolution cryo-EM structure determination in RELION-3. eLife 7, e42166 (2018).

27. Waterhouse, A. et al. SWISS-MODEL: homology modelling of protein structures and complexes. Nucleic Acids Res. 46, W296–W303 (2018).

28. Pettersen, E. F. et al. UCSF Chimera—a visualization system for exploratory research and analysis. J. Comput. Chem. 25, 1605–1612 (2004).

29. Casañal, A., Lohkamp, B. & Emsley, P. Current developments in Coot for macromolecular model building of electron cryo-microscopy and crystallographic data. Protein Sci. 29, 1055–1064 (2020).

30. Liebschner, D. et al. Macromolecular structure determination using X-rays, neutrons and electrons: recent developments in Phenix. Acta Crystallogr. D 75, 861–877 (2019).

31. Croll, T. I. ISOLDE: a physically realistic environment for model building into low-resolution electron-density maps. Acta Crystallogr. D 74, 519–530 (2018).

32. Prisant, M. G., Williams, C. J., Chen, V. B., Richardson, J. S. & Richardson, D. C. New tools in MolProbity validation: CaBLAM for cryoEM backbone, UnDowser to rethink “waters,” and NGL Viewer to recapture online 3D graphics. Protein Sci. 29, 315–329 (2020).

33. Krissinel, E. & Henrick, K. Inference of macromolecular assemblies from crystalline state. J. Mol. Biol. 372, 774–797 (2007).

34. Keuler, T. et al. Development of fluorescent and biotin probes targeting NLRP3. Front. Chem. 9:642273 (2021).

35. Zhang, L. et al. Cryo-EM structure of the activated NAIP2-NLRC4 inflammasome reveals nucleated polymerization. Science 350, 404–409 (2015).

36. Park, K. et al. (2015). Control of repeat-protein curvature by computational protein design. Nat. Struct. Mol. Biol. 22, 167–174 (2015).

